# Individual-Level Traits Outweigh Neighborhood and Landscape Level Factors for Frugivorous Insect Parasites in Small Forest Patches

**DOI:** 10.64898/2026.06.24.734278

**Authors:** Thomas Back, Nolan Miller, Suann Yang

## Abstract

Frugivorous insect larvae are dependent on fruiting plants for development, leading to complex host-parasite interactions that may be influenced by many factors at various scales. We compared the relative effects of factors at the individual, neighborhood, and landscape scales in forest patches. Our results suggest that in areas like upstate New York, where agricultural land uses are dominant, individual scale factors are the most influential. Specifically, parasitism increased with host fruit crop size, but was not associated with host species richness or proximity to forest edge. Notably, the most parasitized hosts were non-native species, including *Frangula alnus* Mill. (Glossy Buckthorn), indicating a potential role of invasive species to shape host-parasite interactions in our system. Our results underscore the importance of host-specific traits in structuring parasitism patterns and suggest management could consider both the ecological context of host traits and the influence of invasive species at multiple scales.

## Introduction

Frugivorous insects that develop within fruits are parasites known to regulate host populations by influencing pre-dispersal seed mortality (Anderson, 1978, Ctvrtecka et al. 2016). While life history traits such as phenology and ontogeny have shaped the evolutionary relationship between insect parasites and plant hosts (Boivin et al. 2015), ecological interactions between them can be influenced by a multitude of factors that occur at different spatial scales. For example, the number of fruits on a given plant (fruit crop size) could influence the way in which hosts are parasitized (e.g. Lee et al. 2001, Pelton et al. 2007, Santoiemma et al. 2017). On the other hand, at the neighborhood scale, local distribution and diversity of hosts within a habitat might affect the ways parasites select hosts (e.g. Inouye 2005). Finally, at the forest patch scale the location of a host on the landscape can impact how the parasites locate hosts (e.g. Basoalto et al. 2010, Buck et al. 2023). While these factors operate at different spatial scales simultaneously, each factor may not have the same level of influence on the host-parasite interaction. Furthermore, the relative influence of factors at different scales may vary with habitat fragmentation, which could disrupt the influence of landscape-scale factors through the loss of habitat connectivity. In the Northeastern United States, around 40% of forests are considered fragmented, commonly by anthropogenic land uses such as agricultural or development (Riitters et al. 2012), underscoring the need to investigate host-parasite interactions in fragmented landscapes.

While the combined impacts of these factors and their relative influence is understudied, evidence from previous studies on the separate effects of these factors offer insight into their combined effects (Basoalto et al. 2010, Inouye 2005, Lee et al. 2015). At the scale of the individual host, factors that determine whether a host is compatible with a parasite are species-specific traits shaped by their coevolutionary history (Boivin et al. 2015). Frugivorous parasites oviposit their eggs on or into the fruits of their host and their larvae develop within the hosts before exiting prior to adulthood (e.g., Basoalto et al. 2010, Elsensohn et al. 2021, Inouye, 2005). For a compatible host, factors that determine whether a host is parasitized include factors such as fruit type size, abundance, ripeness, and phenology. For example, the oviposition site chosen by *Drosophila suzukii* (Matsumura) (Spotted-wing Drosophila) is highest in fruits that are fully ripe and of high nutrient content (Elsensohn et al. 2021). On the other hand, Lepidopterans have been shown to rely on characteristics that they can assess visually (Elsensohn et. al, 2021, Renwick et al. 1994). Higher rates of parasitism can also be associated with larger fruits and more abundant fruit crop sizes (Ctvrtecka et al. 2016). Thus, we would expect to see a relationship between individual host taxonomic groups and occurrence of parasitism, with some host groups that are more prone to parasitism than others because they exhibit the traits preferred by parasites.

On the neighborhood scale, host availability, measured by host species composition and fruit or plant density can have an impact on the occurrence of parasitism. For generalist parasites, species composition surrounding a potential host plant can influence whether the host is parasitized. For example, *Rubus allegheniensis* Porter (Allegheny Blackberry) had lower rates of parasitism where *Prunus serotina* Ehrh. (Black Cherry) was more abundant (Roche et al. 2024). Janzen (1970) and Connell (1970) showed a positive relationship between prey density and rate of predation; thus, higher rates of parasitism should be observed where fruit density is higher, either areas where there are more plants, or where there are plants that have larger fruit crop sizes. The Janzen-Connell hypothesis also applies to the landscape scale where we would expect to see higher rates of parasitism in cultivated areas during the season when fruit crops are highly concentrated (e.g., Basoalto et al. 2010).

At the forest patch scale, heterogeneity can impact the opportunity for interaction between plant host and insect parasite by influencing host availability (Inouye 2005), such as in areas where forest patches adjoin orchards or fields where fruiting plants are cultivated. In the Northeastern U.S. fruiting plants in these forests may share generalist insect parasites with fruit-bearing cultivated plants (e.g., Spotted-wing Drosophila; Roche et al. 2024). When fruits are absent from cultivated areas, alternate hosts within adjacent forests have been found to be incredibly important to frugivorous parasites (Inouye 2005, Wissinger 1997, Tonina et al. 2018). Furthermore, pest management practices can reduce the frequency of parasitism in cultivated plants, though adjacent forests may serve as refuges for insect parasites (Buck et al. 2023), causing a gradient of decreasing parasitism away from forest edges into the cultivated habitat (Tonina et al. 2018). Parasitism at forest edges may be lower than in the forest interior, as avian insectivores are more likely to exploit edge habitats (Lindell et al. 2007). However, in these landscapes, the patterns of parasitism from forest edge to the interior of forest patches needs further study.

In our study we investigated the relative influence of individual, neighborhood, and landscape factors on the occurrence of parasitism in upstate New York. In the agricultural regions of upstate New York, natural forests occur as fragments within a matrix of human-dominated land use. We sampled fruiting plants from forest fragments adjacent to cultivated crop fields, recording the taxonomic group of the host, the species and density of other potential hosts, and the distance from cultivated area to determine the scale of the most influential factor in a fragmented landscape.

### Field Site Description

We obtained samples from seven farm sites across upstate New York (Fig. 1). All sampling occurred in forested areas adjacent to cultivated fields. The average size of forest patches was 5.2 hectares. Sampling locations grew a variety of mostly perennial crops such as blueberries, strawberries and grapes, but also annual crops like corn.

**Figure 1.**
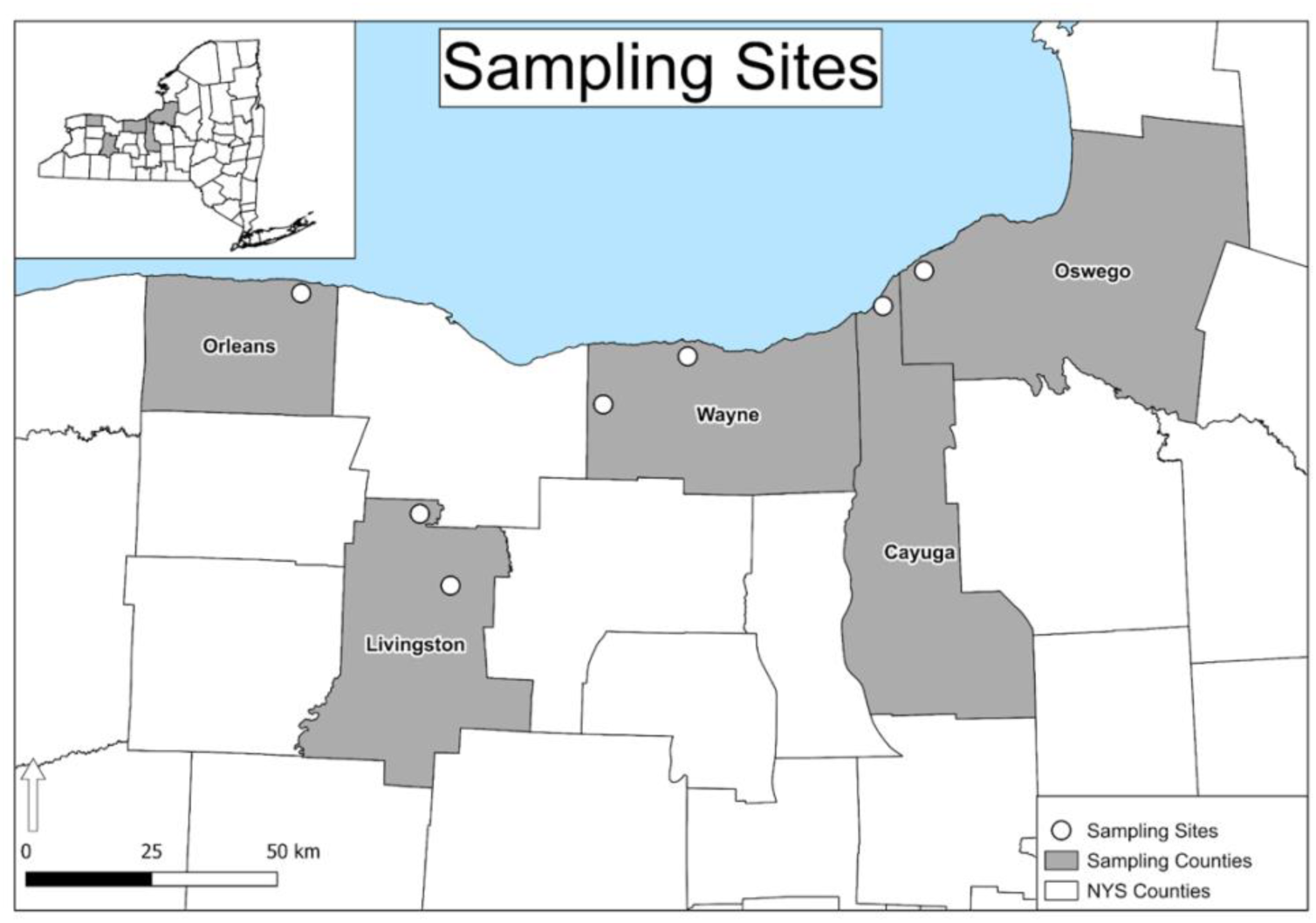
Location of study sites, represented by white circles. We collected fruit from seven farms across five counties in Western and Central New York.

## Methods

### Sampling Design

We sampled fruits for larvae in 2023 and 2024. Two farms were sampled in 2023 between October and November. In the summer of 2024 these two farms were resampled and an additional five farms were added. Each farm was sampled twice, once during early summer (June and early-July) and again during late summer (late-July and August). This timing allowed us to sample while the fruits were ripening and insects were most active (Forest et al. 2010). For each farm, we used QGIS to place sampling plots 250 m apart in a diamond formation (QGIS.org, 2024). In the field, each plot was located; however, some plots were not sampled for logistical reasons, including time constraints, navigational barriers and farmer-issued restrictions. At each plot, each plant with fruits were numbered, identified, and the total number of fruits were estimated. To estimate the fruit counts, we estimated the volume of the plant containing approximately 100 fruits, then estimated what fraction of the plant made up that volume. If there were fewer than 40 fruits present on the plant only half were collected, if there were more than 40 then 20 fruits were collected. All samples were stored in a cooler, then transferred and preserved in a lab fridge at 4.4 °C until processing. Due to time constraints unprocessed fruits were often stored in the fridge upwards of 2 months before dissection or rearing.

### Parasite Detection Methods

Fruit collected in 2023 were all dissected by hand, under a microscope. Parasite occurrence data collected was binary (presence/absence), per fruit. Ripe fruits were always prioritized for processing over unripe fruits due to the parasitism rates being very low in unripe fruits (Lee et. al 2015). Parasitism was determined by either the presence of a larva when the fruit was dissected or an exit hole on the outside of the fruit. Including exit holes allowed us to detect parasitism that occurred prior to the date of collection. However, for samples collected in 2024, we used a different technique to process our much larger sample size more efficiently. This method yielded similar results to dissecting fruits, with an advantage that suitability of the host for developing into an adult can be ascertained (Lee et al. 2015). We incubated samples from 2024 in a Conviron E7 growth chamber with CMP5090 controller, to rear parasitic insects to the stage of emergence from fruit. The growth conditions were set to a 12-hour day/night cycle with daytime temperature of 25 °C and nighttime temperature of 16 °C. To separate samples, we enclosed each plant’s fruits between two mesh sheets inside a plastic container, allowing for larvae to be collected at the bottom of the container once they exited their fruits (Fig. 2). Containers were checked daily, and larvae removed for flash freezing. Parasite data collected using this rearing method is numerical and represents the rate of parasitism per plant, however to match our previous method we converted to presence/absence. Larvae collected were identified to the order level. To merge the data collected from these two different methods, we summarized the 2023 data (collected by dissection) into the number of dissected larvae per plant.

**Figure 2.**
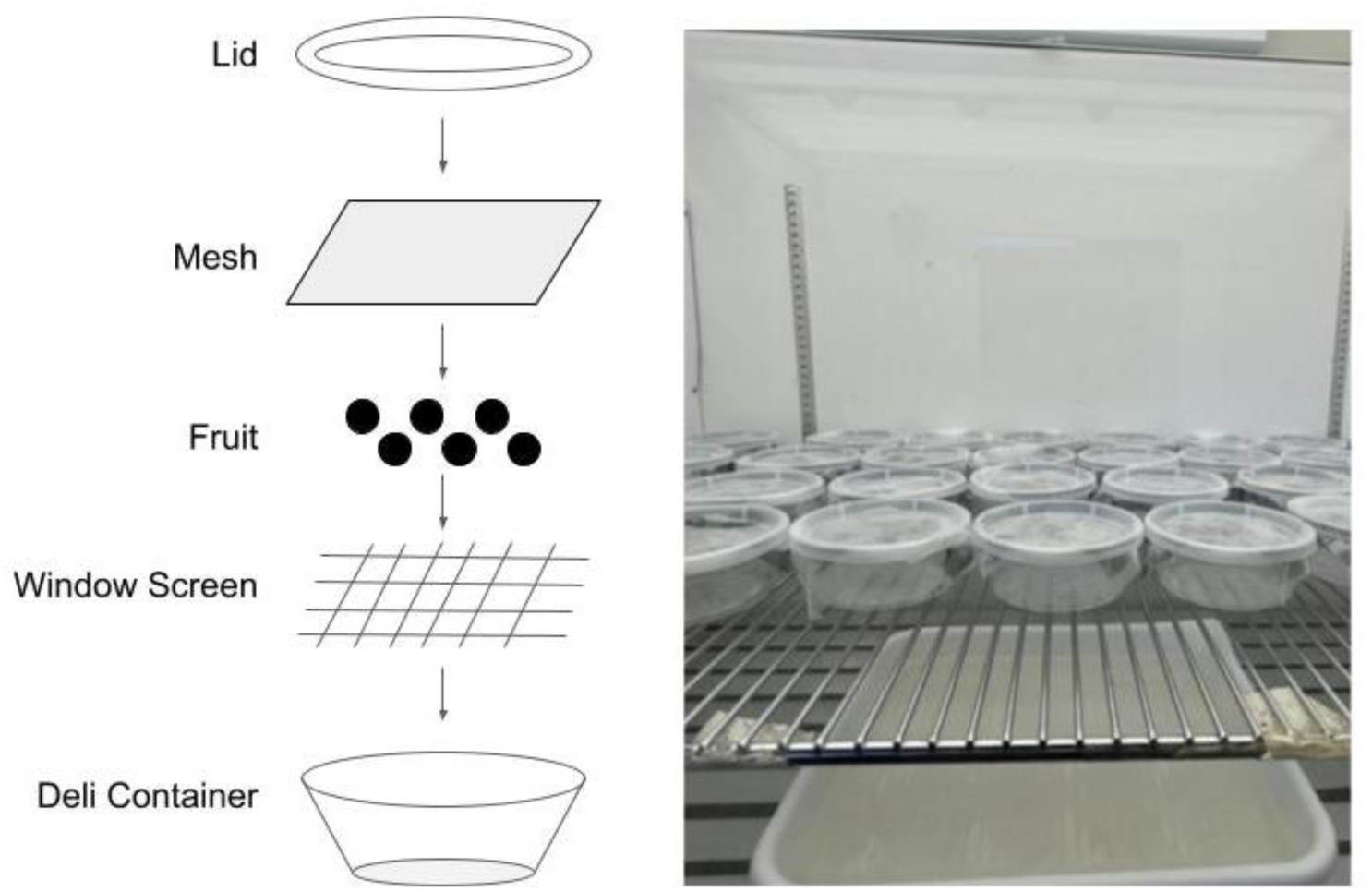
Diagram of larval rearing containers. Each unit contained the fruit from one plant sandwiched between a fine mesh and window screen. This allowed the larvae to escape the fruit and drop into the bottom of the container for easy counting. Adapted from Lee et al. (2015).

### Data Analysis

#### Predictor Variables

The factors investigated at the individual scale were host genus and fruit crop size. The host genus serves as our proxy to understand individual level effects, such as host specific fruit traits (e.g., color, size, water content, etc.). Host genus was used rather than host species due to our limited sample size for host species of low relative abundance. Also at the individual scale, fruit crop size represents how the number of fruits present affects parasitism.

At the neighborhood scale we studied density of potential hosts and species richness at each plot. The density of potential hosts was measured as both the number of plants and number of fruits in a given plot. Species richness was studied to understand the effect of plot level heterogeneity.

We used the distance from the sampling plot to the edge of the forest to investigate the landscape scale. Distance from the plot to the edge of the forest was collected as the minimum distance between the center of the plot and the edge of the forest nearest to the crop field. This distance was calculated using the distance matrix tool inside of QGIS. First, we converted the forest polygon to lines then to points, then obtained the distance between the center of the plot and the nearest point on the edge of the forest. In other words, this is the minimum distance between the sampling plot and the nearest border of the cultivated area.

#### Statistical Analysis

Statistical analyses were conducted with the R Programming Environment (R Core Team 2025). In order to analyze the effects of individual, neighborhood and landscape level factors on the instance of parasitism we used a zero-inflated Poisson mixed effects regression in R using the *glmmTMB* package (Brooks et al. 2017). Because of the potential for correlation between predictor variables at the same scale, we constructed different models consisting of one predictor at each scale (Table 1). We determined the best model by examining AIC values, selecting the models with the smallest AIC to interpret (Table 2).

**Table 1.** Description of explanatory factors used to model host-parasite interactions across scales.

| **Factor** | **Scale** | **Description** | **Used in Models** |
| --- | --- | --- | --- |
| Host Genus | Individual | Identity of host genus, identified in the field | 1, 4, 5 |
| Fruit Crop Size | Individual | Estimated number of fruits on the plant at time of collection | 2, 3, 6 |
| Species Richness | Neighborhood | Number of unique species present in a given plot | 1,2, 3 |
| Plant Density | Neighborhood | Number of individual plants present in a plot | 3, 4 |
| Fruit Density | Neighborhood | Number of fruits present in a given plot | 5, 6 |
| Distance to Edge | Forest Patch | Shortest distance from the edge of the forest to the center of the plot | 1, 2, 3, 4, 5, 6 |

**Table 2.** Model results. Model 2 yielded the lowest AIC, with fruit crop size as the only significant explanatory factor. This highlights the importance of small-scale factors in our system. Models 3-6 not reported due to lack of model convergence.

| Model | Factor | Estimate | $\chi^2$ | df | *P* | AIC |
| --- | --- | --- | --- | --- | --- | --- |
| 1 | Host-Genus | -13.5 | 31.9 | 15 | 0.0066 | 125 |
|  | Species Richness | -0.574 | 2.71 | 1 | 0.0997 |  |
|  | Distance to Edge | -0.0178 | 1.25 | 1 | 0.2640 |  |
| 2 | Fruit Crop Size | 0.00115 | 12.13 | 1 | 0.0005- | 113 |
|  | Species Richness | -0.471 | 1.61 | 1 | 0.1950 |  |
|  | Distance to Edge | 0.0323 | 2.59 | 1 | 0.1080 |  |

## Results

Across our seven farm sites, we were able to collect 29 larvae from 2,638 fruits from 136 plants found in 342 plots. Parasites consisted of members of the insect orders Coleoptera, Lepidoptera and Diptera. Of the sixteen genera sampled larvae were found in only seven (Fig. 3). While the lack of parasitism in rarer genera can likely be attributed to chance, fruit characteristics may explain the lack of parasitism in more common hosts. For example, members of the *Solanum* genus (nightshade) are known for being high in defense compounds (Ben-Abdallah et al. 2019), this could deter hosts. Similarly, we observed that the members of the *Cornus* genus have a notably low pulp volume to seed volume, which would limit the amount of substrate a larva has for development.

**Figure 3.**
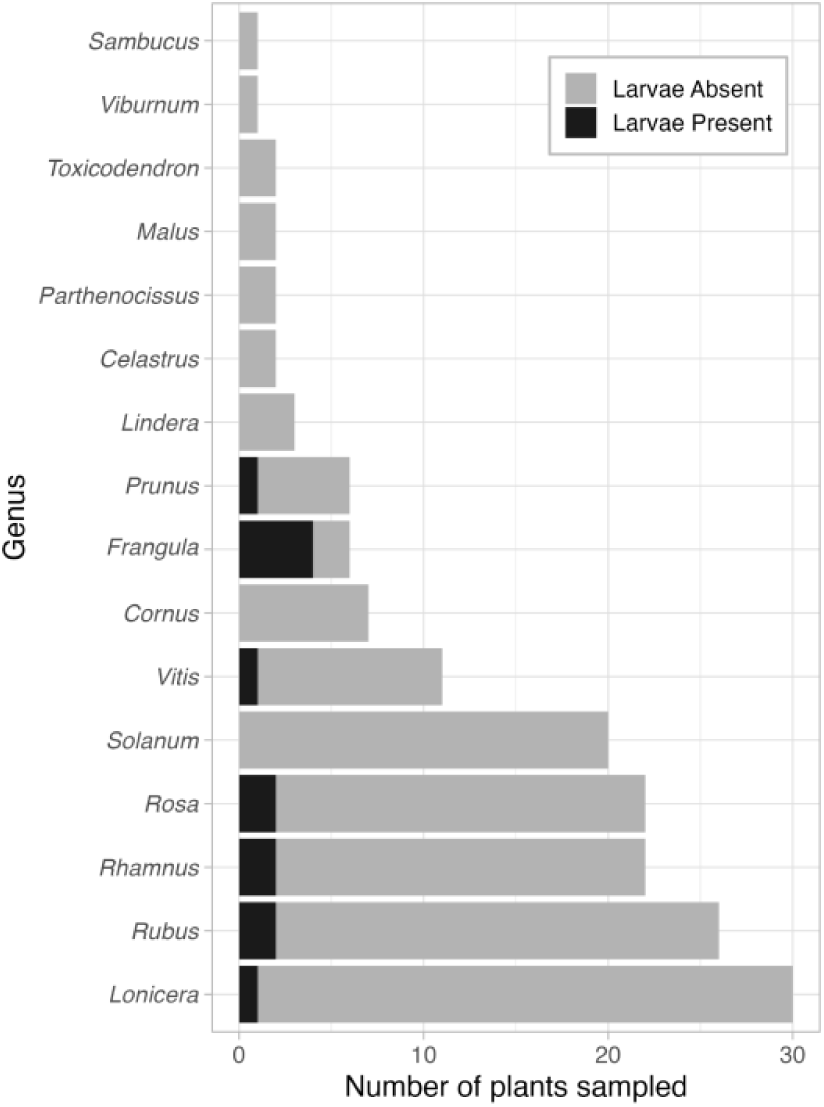
Across the 342 plots, we observed 163 fruiting individuals distributed over 16 genera. Most individuals were not parasitized (gray shading), except for *Frangula* (Glossy Buckthorn). With the exception of *Solanum*, more abundant genera were more likely to have individuals where parasites were observed (black shading).

The best fit model was Model 2 (Table 2; AIC = 113), indicating that plants with a larger fruit crop size had a greater occurrence of parasitism by insect larvae (Fig. 4; Table 2; *P* = 0.000498). The exceptionally high fruit crop size of *Frangula alnus* Mill. (Glossy Buckthorn), likely contributed to this fruit crop size effect, as this species had the highest occurrence of parasites (Fig. 4). At our study sites, Glossy Buckthorn’s median estimated fruit crop size was 1090 fruits per plant, approximately six times more fruits than the next most frequently parasitized host, *Rosa multiflora* Thunb. (Multiflora Rose). We found that all three insect orders (Coleoptera, Diptera, Lepidoptera) were parasites of Glossy Buckthorn. While not a focus of our study, the mechanism behind this high level of parasitism by very different taxonomic groups could be attributed to Glossy Buckthorn’s host potential. Glossy Buckthorn’s large fruit crop sizes and large fruits (approximately fourteen times larger by volume than the next most common host (Multiflora Rose) may offer disproportionately higher quality ovipositing sites to parasites. However, beyond Glossy Buckthorn’s traits related to resource abundance for parasites, host specific traits did not appear to be as influential at the individual level compared to fruit crop size in our system, as host genus was not a predictor in the best fit models (Table 2).

**Figure 4.**
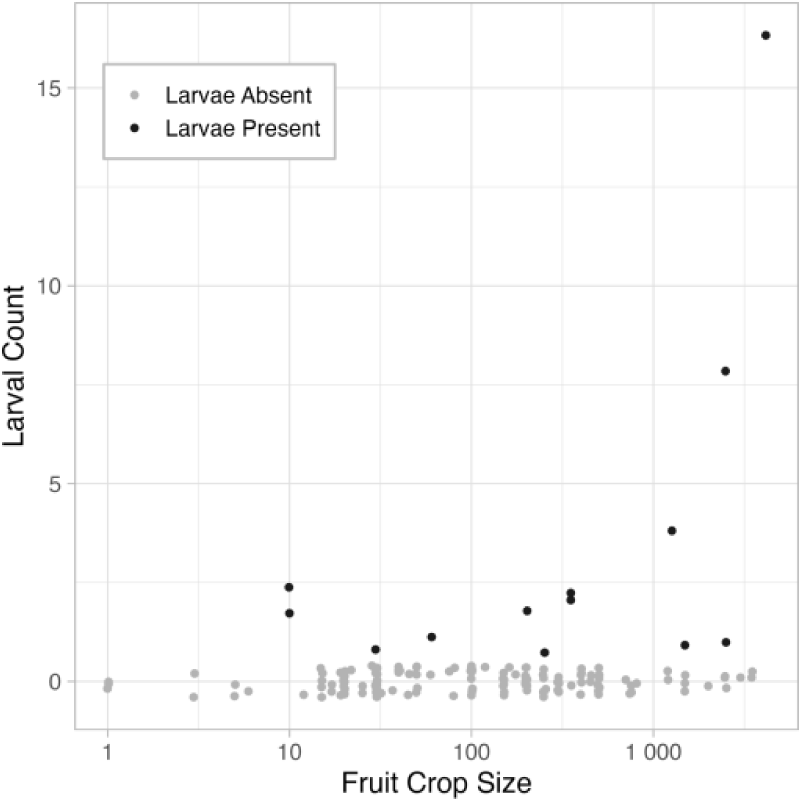
Plants were more likely to be parasitized when fruit crop size was above 10, with higher rates of parasitism as fruit crop size increased (***χ***2 = 14.8, df = 1, *P* = 0.000122). Fruit crop size was the only determining factor for parasitism, suggesting that the individual scale is the most influential level in our system. Note that a small amount of random noise was added to each observation to reduce the overlap of data points.

While individual scale effects (fruit crop size) were influential for occurrence of parasitism, we did not find evidence that plot-level fruiting species richness affects the occurrence of parasitism, suggesting that neighborhood-level effects in our system were not as important as individual-level traits (Fig. 5a; Table 2; *P* = 0.195). However, the lack of relationship between fruiting plant density and the occurrence of parasitism may be attributed to our inability to sample the extremely high plant density areas in our study sites that would be expected to have higher occurrences of parasitism. We encountered thickets of fruiting plant species, such as Multiflora Rose, of such high density that we could not access sampling plots located within them without significant alteration to the forest habitat.

**Figure 5.**
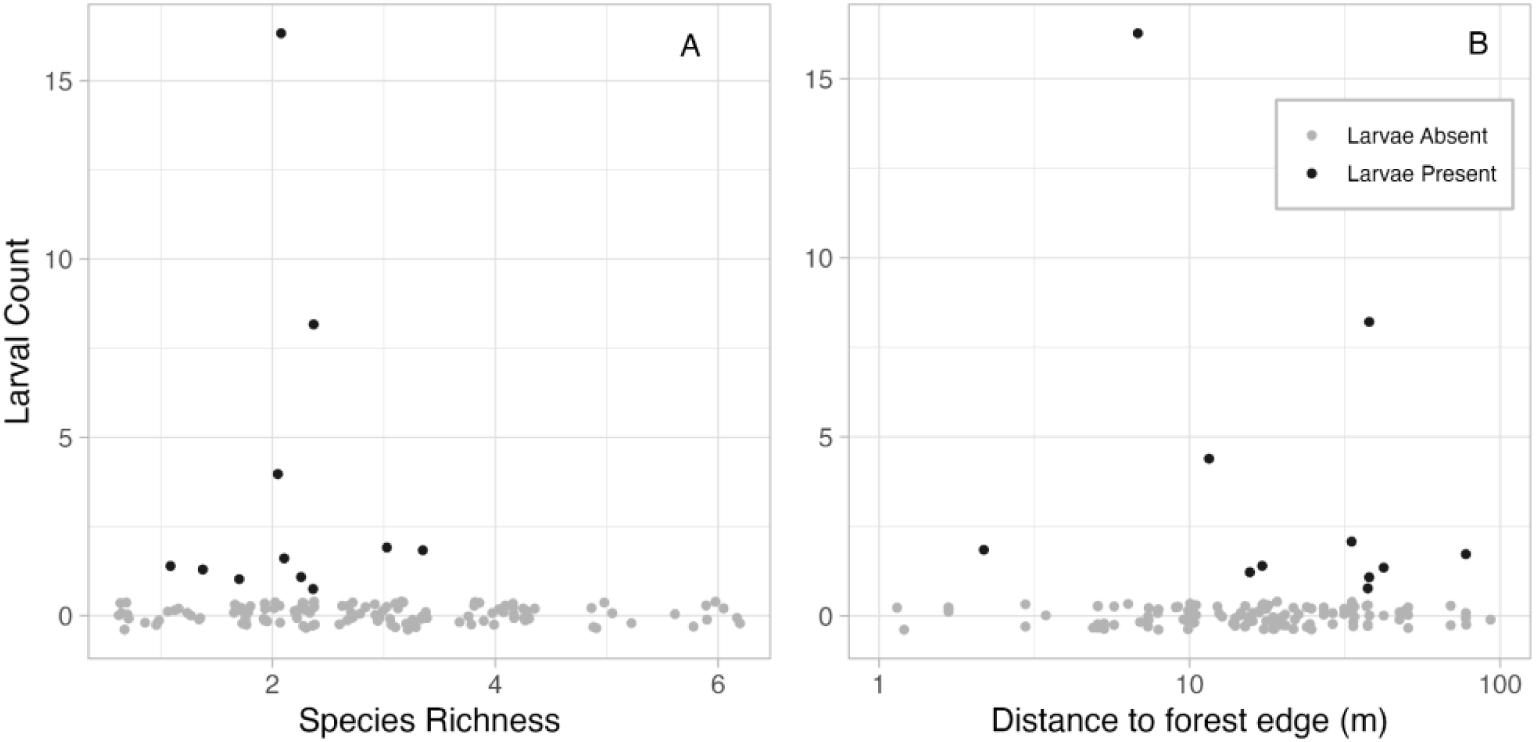
A. The relationship between species richness and the number of larvae found in an individual plant. B. The relationship between forest edge and number of larvae found in an individual plant. Neither variable is a significant factor in the models. Note that a small amount of random noise was added to each observation to reduce the overlap of data points.

Interestingly, while neither plant density nor species richness at the neighborhood level was influential, it is worth noting that of the seven genera that were parasitized, five are not native to upstate New York but were relatively abundant: *Celastrus, Frangula, Rosa, Rhamnus,* and *Lonicera* (Fig. 4). Forest patches surrounded by agriculture and development are more likely to contain invasive species, which could alter host-parasite dynamics (Riitters et al. 2018). While the higher occurrence of parasitism among the non-native hosts could reflect the higher availability of their fruits rather than an innate preference of parasites, the disproportionate parasitism of non-native fruiting species suggests that effective pest management strategies for fruiting crops could include controlling the populations of potential host species in adjacent forests (Buck et al. 2023).

Finally, distance from sampling plot to forest’s edge also did not influence the occurrence of parasitism, indicating that landscape-level effects may also not be as important as individual-level effects (Fig. 5b; Table 2; *P* = 0.108). The lack of a relationship between distance to forest edge and occurrence of parasitism is likely due to the structure of the landscape we studied: a matrix where anthropogenic land uses dominate, with relatively small patches of forest distributed within much larger cultivated or developed habitats. These results imply that, from the perspective of the insects, the entire forest could be considered an edge habitat lacking in forest-interior characteristics. Thus, in systems with small forest patches that lack a gradient from edge to interior habitat, frugivorous parasitic insects may interact with their hosts in a similar way throughout a forest patch.

## Discussion

Our study demonstrates the importance of individual-scale factors on parasitism by frugivorous insect larvae. Unexpectedly, host-specific traits were not as critical as fruit crop size, despite the highly selective nature of oviposition by frugivorous parasitic insects, which often rely on visual and chemical cues to select oviposition sites (Renwick and Chew 1994, Boivin et al. 2012). Our results highlight that in fragmented landscapes, like upstate New York, where agricultural land use is dominant, fruit resource availability at the individual scale may outweigh larger scale factors such as distance to cultivated fields, species richness, and host density.

Our study also provides an example of how the strength of the influence of our neighborhood- and landscape-scale factors on the interaction between insect parasites and their fruiting hosts may depend on forest patch size. Ecological processes operate across scales, and only through studying interactions across scales can we understand the effect of patch size (Wiens 1989, Tscharntke et al. 2012). Our study sites, in a region dominated by agricultural and residential land uses, consisted of very small forest patches (5.2 hectares on average), in contrast to the contiguous forest studied by Roche et al. (2024), where neighborhood effects were detected. Landscapes dominated by human land use typically are characterized by reduced forest land cover, along with associated losses of ecosystem services (Hasan 2020), such as top-down control by predators and the protective effects of species biodiversity. While the effects of habitat fragmentation can be highly species- and system-specific (Debinski and Holt 2000), we may expect to find that individual-level traits are the most influential for interactions between fruiting plants and their parasites in highly fragmented landscapes in general.

In addition, studying these interactions at various scales, including the neighborhood scale, allows us to gain a better understanding of the way invasive species are affecting host-parasite interactions. Non-native plants may serve as more attractive hosts, especially to non-native parasites. In areas where both non-native plants and insects are establishing, the interactions between co-occurring non-native species should be similar to their interactions in their native ranges (Wan et al. 2018). This reassociation may be occurring for the host-parasite interactions we observed. Five genera of the seven that contained parasites were not native to upstate New York (Fig. 4), indicating that most of the host-parasite interactions that were observed involve non-native plants and highlighting the potential impacts of these invasive species beyond competition with other plants. For example, while Dipterans were the least common parasite we found, glossy buckthorn is a viable host for Spotted-wing Drosophila in both the United States and in glossy buckthorn’s native range of Eastern Europe, Western Asia and, Northern Africa (Greenleaf et al. 2023, Grassi et al. 2011). While beyond the scope of our study, greater taxonomic resolution on the larvae found in fruits would allow for questions to be asked about reassociation and novel interactions between invasive plant species and parasites of their fruits.

We suggest that the dominance of invasive plants participating in these host-parasite interactions has management implications. Land managers who are interested in managing parasites may wish to consider reducing the populations of invasive fruiting plants within forests adjacent to cultivated fruiting crops. In our study, the invasive Glossy Buckthorn was more likely to have a parasite than not. Numerous pest insects are associated with Glossy Buckthorn (Greenleaf et al. 2023); thus, the long fruiting phenology of Glossy Buckthorn means this plant species can provide refuge habitat to pest insects when cultivated fruits are out of season or after pesticide application in the cultivated areas. Thus, forests that contain invasive fruiting plant species can serve as unexpected sources of insect parasites for adjacent crop fields. In predictably ephemeral agricultural habitats, these dynamics can have serious economic and ecological consequences (Wissinger 1997).

In summary, in areas dominated by agricultural land uses, we found that individual-scale, resource availability traits were the most influential on the occurrence of parasitism, with the invasive Glossy Buckthorn parasitized at the highest rate. Fragmented landscapes that contain very small habitat patches may have reduced influence from larger-scale factors on the type of species interaction that we studied. Finally, these results are applicable to pest management, where managing invasive species in the forests surrounding the cultivated areas could lead to lower parasites within the cultivated field. Overall, studying host-parasite interactions at multiple scales may be a critical step to identify the scale to target for developing pest management strategies in specific ecosystems.

## Acknowledgments

We would like to thank the Geneseo Foundation for the funding and the Yang and Hoven research groups who provided us with invaluable feedback. We would also like to acknowledge and thank each individual landowner for allowing us to conduct our research on their properties. Finally. we would like to acknowledge and express gratitude to the Haudenosaunee Nation, whose traditional homeland contains all of our sampling sites.

## Notes

### Competing Interest Statement

The authors have declared no competing interest.

